# Machine Learning Speeding Up the Development of Portfolio of New Crop Varieties to Adapt to and Mitigate Climate Change

**DOI:** 10.1101/2021.10.06.463347

**Authors:** A. Bari, H. Ouabbou, A. Jilal, H. Khazaei, F.L. Stoddard, M.J. Sillanpää

**Affiliations:** Operational AI, Montreal, Canada; INRA, Rabat, Morocco; World Vegetable Centre, Tainan, Taiwan; University of Helsinki, Helsinki, Finland; University of Oulu, Oulu, Finland

## Abstract

Climate change poses serious challenges to achieving food security in a time of a need to produce more food to keep up with the world’s increasing demand for food. There is an urgent need to speed up the development of new high yielding varieties with traits of adaptation and mitigation to climate change. Mathematical approaches, including ML approaches, have been used to search for such traits, leading to unprecedented results as some of the traits, including heat traits that have been long sought-for, have been found within a short period of time.

## 1. Introduction

Climate change poses serious challenges to achieving food security. It is a dual challenge that requires keeping up with the world’s increasing demand for food, while both adapting to and mitigating climate change. The agriculture sector is in the midst of climate change, in a time of a need to keep pace and produce more to close the gap of 56% between the amount of food available today and that required by 2050 (World Resources Institute, 2021, Figure 1).

**Figure 1.**
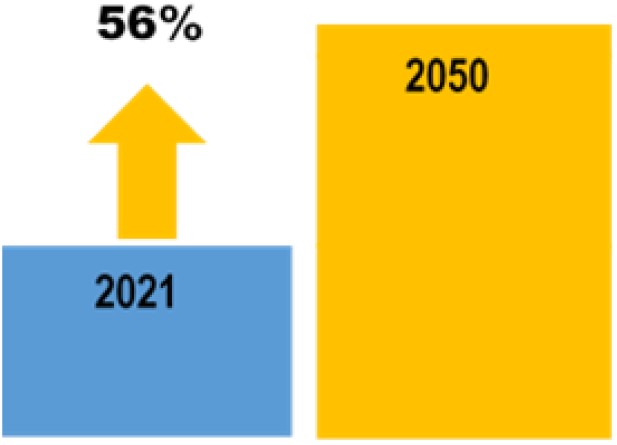
Required Increase to meet the food demand by the year 2050.

To help speed up the development of new crops varieties with traits to adapt and mitigate changing climate conditions, we have used mathematical approaches, including machine learning techniques. Mathematical approaches have played a major role in producing more food, prior to the emergence of molecular approaches. A century ago, Fisher (1930) elaborated the mathematical theoretical framework that was considered as the basis of quantitative genetic theory, on which crop improvement was established, to produce more food. These mathematical approaches have been used in crop improvement to capture genetic variability and inheritance of quantitative traits, assess the interaction between varieties and environments and predict the performance gains in yields (Scheffé, 1959). In their mathematical model, Koo and Wright (2000) found that early identification of valuable crop traits, is of equal importance to the process of transferring these traits into an improved genetic background.

Interest is currently returning to the use of mathematical as well as machine learning approaches to accelerate and optimize further crop improvement, in particular to address networked genes involved in trait expression and also to optimize crop improvement processes by shortening the time and reducing the costs to develop new and portfolio of crop varieties with enhanced traits within a short period of time (Anderssen and Edwards, 2012; Bari et al., 2016; Parmley et al., 2019; Tong and Nikoloski, 2021). There is particular potential for using machine learning to identify economically important traits in genetic resources that can then be incorporated into crop improvement programs that develop portfolios of cultivars for diversified systems. Diversified systems have been reported to raise productivity and improve livelihoods, performing particularly well under environmental stress and delivering production increases in places where additional food is desperately needed (De Schutter and Frison, 2017).

## 2. Methodology

We used different machine-learning techniques to accelerate the search for adaptive traits to drought and heat in crops. The machine-learning technique used span supervised and unsupervised techniques, including Bayes, Neural Network (NN), Random Forest and K-means techniques, to help in the rapid identification and location of adaptive traits (Cherkassky and Mulier, 2007; Khazaei et al., 2013; Bari et al., 2016). The ML-based search for these adaptive traits is based on exploring and exploiting the dependence between the desired traits (denoted Y) and the environment (denoted X) as an evolutionary co-driver prevailing in the areas where the samples were collected (Henry and Nevo, 2014). The desired traits can be considered as representative variables with the additive influence of many genes with small effects (Brown et al., 1996). If this dependence exists, it should be possible to predict the values that could be assigned *in silico* to the unknown samples of crops of those lacking evaluation for these traits. A general regionalized variable model (GRVM) was also used, based on predicted probabilities of the Bayes model to create the map of climatic change performance of the models. Crop simulation models combined with high-resolution climate change map scenarios can help to identify key traits that are important under drought and high temperature stress in crops (Semenov and Halford, 2009).

The environmental data (X) are long-term climate data of the original collection areas where the samples of crops are considered to have evolved. Two types of climate data were used, namely the CGIAR climate surface data and the world climate surfaces (Hijmans et al., 2005). The CGIAR surfaces were generated from meteorological-station data based on the ‘thin-plate-smoothing spline’ method (Hutchinson, 2000). The generation of surfaces included the use of terrain variables as auxiliary variables that were first converted into digital elevation models (DEM) to increase the precision of the interpolated values. The Worldclim data are also gridded data, generated through interpolation of average monthly climate data from global networks of weather stations using the thin-plate smoothing spline procedure. All these climate datasets were of approximately of 1 km square resolution.

The accuracy of all models was measured using values derived from a confusion-matrix table and the Area Under the Curve (AUC) of the Receiver Operating Characteristics (ROC) (Swets et al., 2000; Fawcett, 2006). The Kappa statistic was used also to measure the specific agreement between predictions and observations in the confusion matrix table. High values of both AUC and Kappa indicate that the model’s performance is adequate for prediction purposes (Scott et al., 2002).

### 2.1 Adaptation to drought

To identify traits of adaptation to drought, we used the faba bean (*Vicia faba* L.), which is a widely grown food legume crop in the dry areas and thus considered as one of the most likely crops to be impacted by climate change (Duc et al., 2011). Cumulative plant datasets from experiments on faba bean are used to explore the link between the trait expression (Y) and the environment (X). A total of 400 plant samples was originally selected on the basis of extreme “wet” and “dry” environmental profiles using a clustering algorithm from 10,000 samples stored in CGIAR genebanks. From these 400, a first subset was used to detect the presence of patterns, if any, in the data form the training set, and the patterns or dependence detected together with a new X was then used in turn to assign values, as predictive probabilities of having the sought-after drought traits, to another unknown test set. The plants of this latter subset were grown for evaluation, beside the first set, to test the predictions for their accuracy and agreement with the actual evaluation data. The trait plant data consists of a set of leaf morpho-physiological measurements that capture drought-adaptation related traits used previously (Khazaei et al., 2013). These measurements span plant gas exchange properties, photosynthesis and phenology. The assertion of the presence and absence of the traits of drought tolerance in the samples was tested and validated (Khazaei et al., 2013).

### 2.2 Adaptation to heat

Here we used unsupervised learning to cluster data, barley (*Hordeum vulgare* L.) was the test crop plant species. Clustering was conducted using environmental data (*a priori* information) to develop one subset that was likely to contain climate-change–related traits and another subset representative of the different environments where the barley samples were originally sampled across Morocco. The samples allocated to any one cluster shared phenotypic affinity vis a vis either presence or absence of the trait of tolerance to heat. Each subset contained 100 samples, of which 30 samples were selected at random. Another core subset was formed to capture most of the diversity, of which also 30 were drawn at random (Figure 2). All the subsets were grown in the same field for comparison based on *a posteriori* evaluation. The environmental data used in the partitioning consisted of 19 climate variables extracted from world climate data.

**Figure 2.**
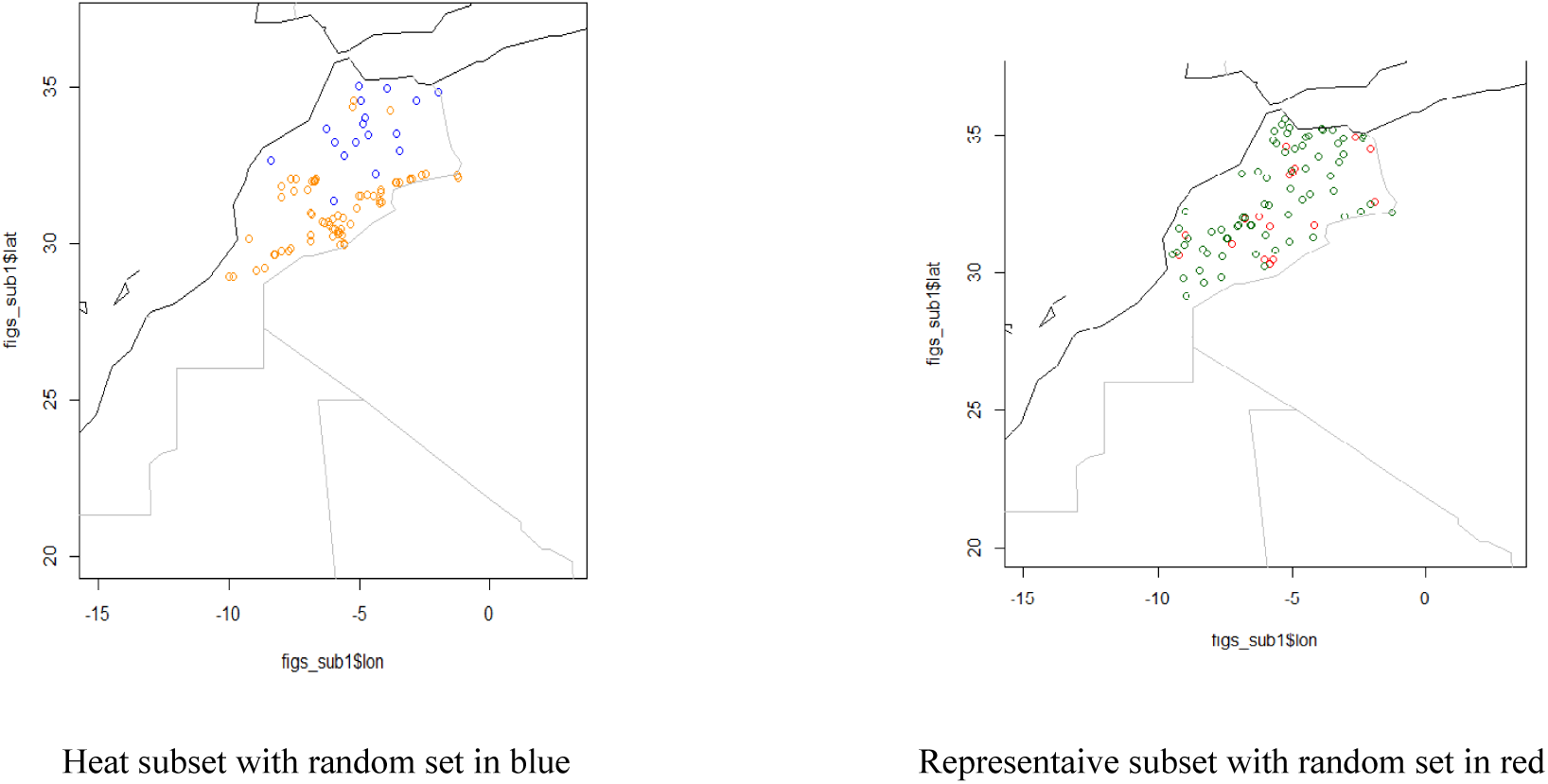
Sites of samples of barley of both subsets with targeted subset (left) and representative subset (right).

This part of the partitioning process is to identify traits that breeders have long sought-for in order to combine an optimized grain-filling period with maturity rather than with earliness alone. Partitioning has been carried out for both the core subset and the trait-based subsets, in particular, to assess plant genetic resources of barley for heat traits based on agro-morphological data and climate data.

To compare and validate the sub-setting based on climate data, the samples were compared on their evaluation attributes. Both subsets were grown among 697 entries at the Moroccan INRA experimental station (33.605°N; 06.716°W; 410 m altitude) in single rows of 5 m length. Observations were taken on several agro-morphological and physiological traits. After harvesting, all the samples were passed through the Near-infrared spectroscopy (Infraneo machine) and the absorbance data (from 850 nm to 1048 nm) with protein content were extracted. Further evaluation was carried out using a honeycomb design of hill plots with the aim to also reduce the area needed for evaluation. Subsequent evaluation was also carried out focusing mostly on the two subsets and heat-related traits, among which were canopy temperature and phenology traits, specifically grain-filling period.

## 3. Results and discussion

### 3.1 Drought adaptation

There was strong agreement between the prediction and the actual evaluation of the plants in terms of their capability to tolerate drought. Both AUC and Kappa values were high, indicating that it is highly likely to identify traits that will provide stress tolerance to crops and can be transferred to cultivars by breeding (Table 1).

**Table1.** Performance metric values (case of NN model).

| Subset |  | Training set |  |  | Test set |  |  |
| --- | --- | --- | --- | --- | --- | --- | --- |
| Statistic | AUC | %Correct | Kappa |  | AUC | %Correct | Kappa |
| Mean | 0.96 | 0.96 | 0.92 | Mean | 0.88 | 0.88 | 0.77 |
| SD | 0.02 | 0.01 | 0.03 | SD | 0.04 | 0.04 | 0.08 |
| CI 95% | ±0.01 | ±0.01 | ±0.02 | CI 95% | ±0.02 | ±0.02 | ±0.05 |

The results show (Figure 3) that machine learning can help in reducing field evaluation, as testing can be focused on those samples that are highly likely to have the desired traits rather than having to screen the whole collection, which is practically impossible, given the large number of samples. This thus will reduce costs, particularly for evaluation of the so many samples that are not likely to have the desired traits. This can help also in speeding up the development of new crop varieties, while minimising costs.

**Figure 3.**
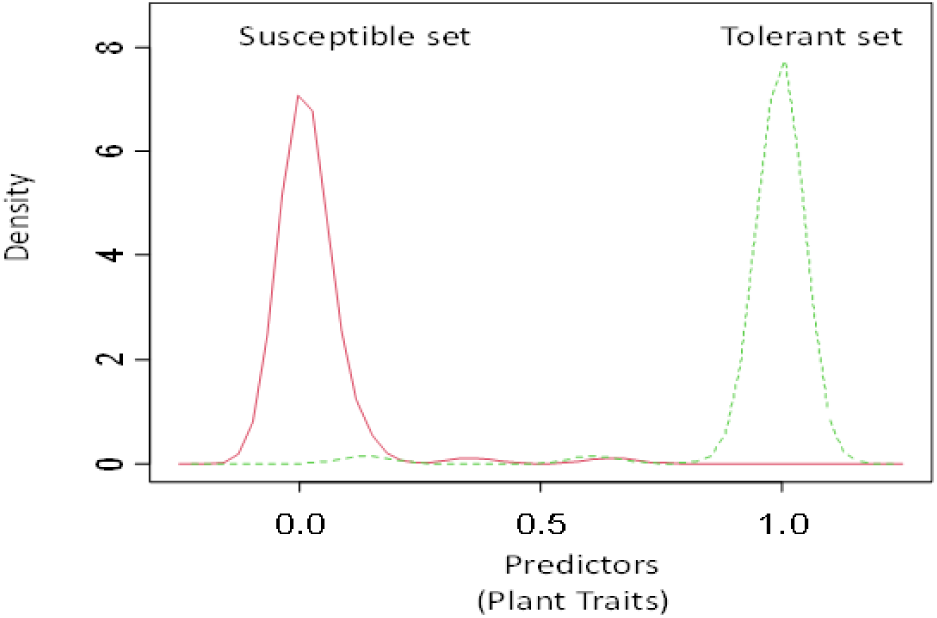
Density plots of prediction for tolerance and susceptible samples for the ML model using test set; the green (broken) curve indicates the probability density distribution for adaptation to drought and the red (solid) curve indicates susceptibility to drought.

### 3.2 Heat adaptation

In terms of adaptation to heat, the results show that it is possible to identify the sought-after heat traits even with unsupervised learning prior to evaluation. The a posteriori evaluation involving heat-related traits such as days to maturity (a measure of phenology) and canopy temperature, especially in the grain-filling period, showed that the two subsets were different. The heat subset tended to have lower leaf temperature than the core or random subsets (Figure 4), so it tolerated the heat better and the heat subset is thus more likely to yield heat-related traits. These results indicates that natural genetic variation contains the much-needed trait variation to help with heat tolerance in crop improvement.

**Figure 4.**
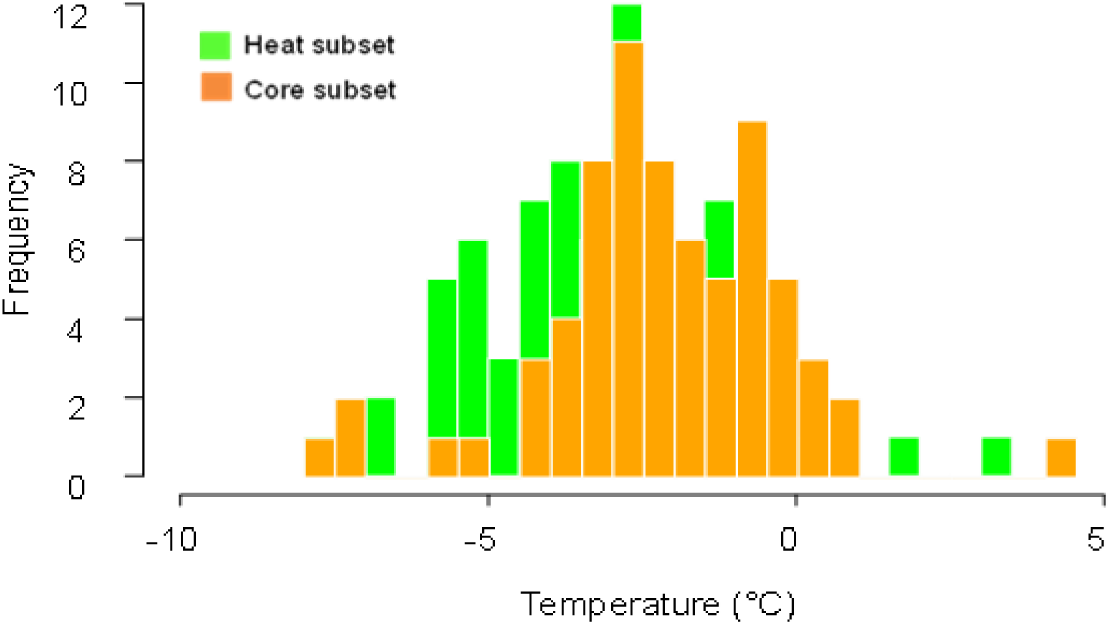
Canopy temperature depression of the heat subset vs the core subset (representative set). The heat subset lowers its temperature when compared to core and random subsets. A trait that helps to withstand heat.

## 4. Conclusions

Mathematical models including machine-learning models can help tremendously in identifying the desired traits while shortening the time and potentially reducing significantly the costs to develop new crops. Machine learning can help to reduce costs that are incurred to assess and evaluate large number of samples in genebanks. There are more than 1,750 genebanks worldwide, holding together more than 7 million plant samples. ML has the potential to identify rapidly traits, including climate-related traits, and to speed up the crop development processes to develop portfolios of crops varieties to adapt to and mitigate climate change.

## References

Anderssen RS, Edwards MP (2012) Mathematical modelling in the science and technology of plant breeding. Int J Numer Anal Model Series B 3:242–258.

Bari A, Khazaei H, Stoddard FL, Street K, Sillanpää MJ, Chaubey YP, Dayanandan S, Endresen DF, De Pauw E, Damania AB (2016) In silico evaluation of plant genetic resources to search for traits for adaptation to climate change. Clim Change 134(4):667–680. http://dx.doi.org/10.1007/s10584-015-1541-9

Bari A, Damania AD, Mackay M, Dayanandan S (2016) Applied Mathematics and Omics to Assess Crop Genetic Resources for Climate Change Adaptive Traits. CRC Press, Taylor & Francis Group, Boca Raton, FL, USA. https://www.routledge.com/products/9781498730136

Brown JH, George Stevens GC, Kaufman DM (1996) The geographic range: size, shape, boundaries, and internal structure. Annu Rev Ecol Syst 27:597–623. https://doi.org/10.1146/annurev.ecolsys.27.1.597

Cherkassky V, Mulier F (2007) Learning from Data: Concepts, Theory, and Methods. Wiley-IEEE Press. 560 p. ISBN:0471681822

De Schutter O, Frison E (2017) Modern agriculture cultivates climate change – we must nurture biodiversity. Grain. https://grain.org/es/article/entries/5634-modern-agriculture-cultivates-climate-change-we-must-nurture-biodiversity

Duc G, Link W, Marget P, Redden RJ, Stoddard FL, Torres AM, Cubero JI (2011) Genetic adjustment to changing climates: Faba bean. In: Yadav SS, Redden RJ, Hatfield JL, Lotze-Campen H, Hall AE (Eds.), Crop adaption to climate change, 1rd ed. John Wiley & Sons, pp. 269–286.

Fawcett T (2006) An introduction to ROC analysis. Pattern Recogn Lett 27:861–874.

Fisher R (1930) Inverse Probability. Mathematical Proceedings of the Cambridge Philosophical Society 26(4):528–535. http://dx.doi.org/10.1017/S0305004100016297

Haldane JBS (1924) A mathematical theory of natural and artificial selection. Part I. Transactions of the Cambridge Philosophical Society 23:19–41.

Henry RJ, Nevo E (2014) Exploring natural selection to guide breeding for agriculture. Plant Biotechnol J 12:655–662. https://doi.org/10.1111/pbi.12215

Hijmans RJ, Cameron SE, Parra JL, Jones PG, Jarvis A (2005) Very high resolution interpolated climate surfaces for global land areas. Int J Climatol 25:1965–1978. https://doi.org/10.1002/joc.1276

Hutchinson MF (2000) ANUSPLIN Version 4.1 User’s Guide. Australian National University, Centre for Resource and Environmental Studies, Canberra.

Khazaei H, Street K, Bari A, Mackay M, Stoddard, FL (2013) The FIGS (focused identification of germplasm strategy) approach identifies traits related to drought adaptation in Vicia faba genetic resources. PLoS One 8(5):e63107. https://doi.org/10.1371/journal.pone.0063107

Koo B, Wright BD (2000) The optimal timing of evaluation of genebank accessions and the effects of biotechnology. Am J Agric Econ 82:797–811. https://www.jstor.org/stable/1244521

Parmley KA, Higgins RH, Ganapathysubramanian B, Sarkar S, Singh AK (2019) Machine learning approach for prescriptive plant breeding. Sci Rep 9:17132. https://doi.org/10.1038/s41598-019-53451-4

Poggio T, Rifkin R, Mukherjee S, Niyogi P (2004) General conditions for predictivity in learning theory. Nature 428:419–422. https://doi.org/10.1038/nature02341

Scheffé H (1959) The analysis of variance. Wiley, New York. 477 p. https://doi.org/10.1002/bimj.19610030206

Semenov MA, Halford NG (2009) Identifying target traits and molecular mechanisms for wheat breeding under a changing climate. J Exp Bot 60:2791–2804. https://doi.org/10.1093/jxb/erp164

Swets JA, Dawes RM, Monahan J (2000) Better decisions through science. Sci Amer 283(4):82–87. https://www.jstor.org/stable/26058901

Tong H, Nikoloski Z (2021) Machine learning approaches for crop improvement: Leveraging phenotypic and genotypic big data. J Plant Physiol. 257:153–354. https://doi.org/10.1016/j.jplph.2020.153354

World Resources Institute (2021). Creating a Sustainable Food Future. https://www.wri.org/research/creating-sustainable-food-future

